# Resolving out of Africa event for Papua New Guinean population using neural network

**DOI:** 10.1101/2024.09.19.613861

**Authors:** Mayukh Mondal, Mathilde André, Ajai K. Pathak, Nicolas Brucato, François-Xavier Ricaut, Mait Metspalu, Anders Eriksson

## Abstract

The demographic history of the Papua New Guinean population is a subject of significant interest due to its early settlement in New Guinea, at least 50 thousand years ago, and its relative isolation compared to other out of Africa populations. This isolation, combined with substantial Denisovan ancestry, contributes to the unique genetic makeup of the Papua New Guinean population. Previous research suggested the possibility of admixture with an early diverged modern human population, but the extent of this contribution remains debated. This study re-examines the demographic history of the Papua New Guinean population using newly published samples and advanced analytical methods. Our findings demonstrate that the observed shifts in relative cross coalescent rate curves are unlikely to result from technical artefacts or contributions from an earlier out of Africa population. Instead, they are likely due to a significant bottleneck and slower population growth rate within the Papua New Guinean population. Our analysis positions the Papua New Guinean population as a sister group to other Asian populations, challenging the notion of Papua New Guinean as an outgroup to both European and Asian populations. This study provides new insights into the complex demographic history of the Papua New Guinean population and underscores the importance of considering population-specific demographic events in interpreting relative cross coalescent rate curves.

## Introduction

The Papua New Guinean (PNG) population is among the most fascinating in the world, owing to its unique demographic history. Following the Out Of Africa (OOA) event, modern humans populated New Guinea at a remarkably early date-at least 50 thousand years ago^1^. Since then, the population has remained relatively isolated compared to other OOA populations (such as European and Asian populations)^2–5^ and has gone through a strong bottleneck ^6^. The substantial Denisovan ancestry within the PNG population^7,8^ and the strong correlation between Denisovan and Papuan ancestries^9^, contribute to the genetic distinctiveness of the PNG population.

Researchers have suggested that the genomes of PNG populations contain evidence of admixture with a modern human population that might have diverged from African populations – around 120 thousand years ago – much earlier than the proclaimed primary divergence between African and OOA populations^4,10^. However, the extent to which this early diverged population contributed to the genome of PNG populations remains a subject of ongoing debate^3,4,11,12^. Interestingly, this early migration hypothesis is more widely accepted by archeologists^13–17^.

Pagani et al^4^ supports this hypothesis, notably through Relative Cross-Coalescent Rate (RCCR) analysis. This RCCR analysis suggests that the PNG population diverged from African populations significantly earlier than other OOA populations. They argued that this earlier divergence indicated by the RCCR curve might reflect a contribution from an earlier OOA population specific to PNG. While this shift in the RCCR curve is well-documented^2,3,5^, some researchers attribute it to technical artefacts such as low sample sizes and phasing errors rather than genuine demographic events.

The origins of the primary lineage of the PNG population have also been contested. Some researchers propose that the PNG population is closely related to the Asia-Pacific populations and serves as a sister group to other Asian populations^11,12,18^. Conversely, other researchers argue that the PNG population is an outgroup to both European and Asian populations^3,9,19^. Recent advancements in analytical methods may provide new insights into these debates. For example, Approximate Bayesian Computation with Deep learning and sequential Monte Carlo (ABC-DLS) allows for the use of any summary statistics derived from simulations to train neural networks, which can then predict the most likely demographic models and parameters based on empirical data^20^. Additionally, the Relate software^21^ enhances RCCR analysis by employing a modified version of the hidden Markov model, initially used in the Multiple Sequentially Markovian Coalescent (MSMC) method^22,23^, allowing for the analysis of thousands of individuals with greater robustness.

In this paper, we re-examine the demographic history of the PNG population using newly published samples^24^ combined with data from the 1000 Genome Project^25,26^ and cutting edge methods^20,21^. This approach has enabled us to address these longstanding questions with greater precision. We first generate new empirical RCCR curves and demonstrate that the previously observed shift is unlikely to be the result of low sample size or phasing errors. Through simulations, we further show that the PNG population is indeed a sister group to other Asian populations and this shift is probably not due to contributions from an earlier OOA population. Instead, it is likely a consequence of a significant bottleneck and slower population growth in the PNG population.

## Results

### Analysis of empirical data

We used Relate^21^ to estimate the effective population size and track RCCR changes over time. To ensure robustness, we performed this analysis using 40 random samples per population, repeated 10 times to generate confidence intervals (Figure 1). The results for effective population size (Supplementary Figure 1) align closely with previous findings^2,5,21^. However, RCCR analysis reveals a significant earlier divergence between the PNG highlander population and African population (using Yoruba as the reference) around 100,000 years ago, with a RCCR value of 68% (66-70%). This contrasts with other OOA populations such as Europeans (83% [81-85%]) and East Asians (83% [82-84%]) at the same time point. A similar shift was also found when lowlanders from PNG were used (Supplementary Figure 2).

**Figure 1:**
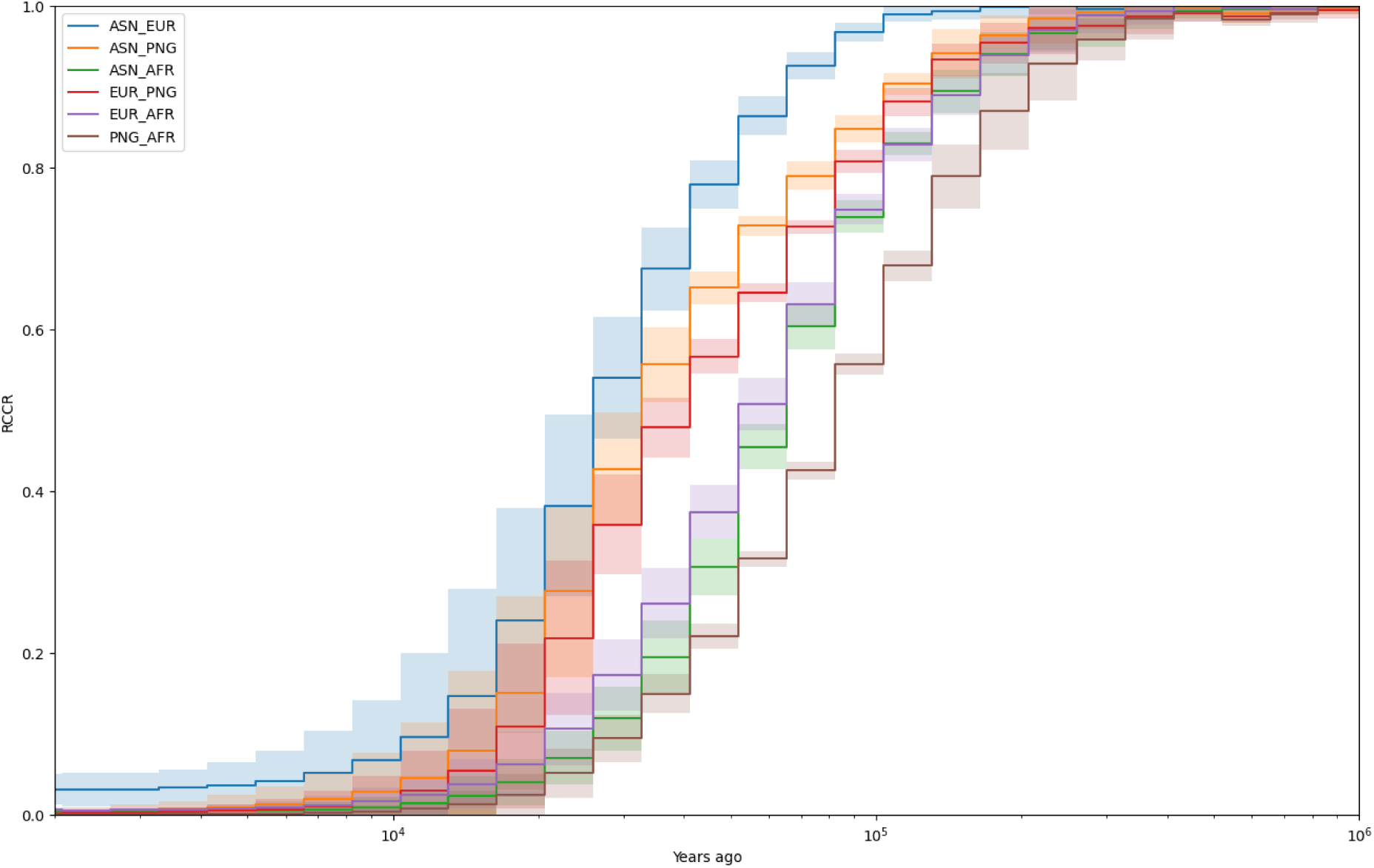
Relate results of relative cross coalescent rate (RCCR) curve on empirical data using 40 samples per population. AFR = Yoruba, EUR = British from England and Scotland, ASN = Han Chinese and PNG = Highlanders of Papua New Guinea. The x-axis is years ago from modern times with the log scale. The y-axis is RCCR values calculated by Relate. The shaded regions are the 95 percent (two standard deviation) confidence intervals created from 10 separate random sampling events. The line represents the mean values.

This earlier separation in PNG has been noted by various studies using MSMC analyses^2,4,5,22^. Pagani et al.^4^ suggested that this shift might be due to a contribution from the first OOA population to the PNG, while others have speculated that it could be due to phasing errors in the PNG genome^2,27^. Given that we phased the genomes using 249 samples from PNG (Supplementary Table 7, see André et al.^24^ for more details about the populations), the likelihood of systematic phasing errors causing this shift is minimal^28^ and are therefore an unlikely explanation for this shift as claimed before^2,27^. This conclusion is further supported by analyses using physically mapped datasets (Supplementary Figure 3) from the SGDP^27^, although these results are less definitive due to the smaller sample size (n=2 per population). A similar shift (Supplementary Figure 4) was also noted in the Andamanese population^12^ though the sample size for Andamanese is smaller as well (n=10). The Andamanese are particularly interesting as they do not have sizable Denisovan ancestry or contributions from an earlier OOA population^11,12,18^, which helps to narrow down the possible causes for the RCCR shift.

### Reconstructing demographic history of PNG

To explore the demographic processes causing the observed RCCR shift, we tested five plausible demographic scenarios labelled A, O, M, AX and OX (Supplementary Figure 5). In Model A, the PNG and East Asian populations are sister groups^11,12,18^. Model O positions the PNG population as an outgroup to both European and East Asian populations^3,19^. Model M combines elements of both A and O, suggesting that the PNG population arose from admixture between a sister group of the Asian population and an outgroup of European and Asian populations. Model AX postulates that the PNG population is a sister group to the Asian population but received input from an earlier OOA population^4^. Finally, in Model OX, the PNG population receives a contribution from an earlier OOA population, while remaining ancestry came from an out group to the European and East Asian populations (see methods for further details).

We trained our neural network using cross-population Site Frequency Spectrum (cSFS) data under these five simulated models (Supplementary Table 1, Supplementary Figure 5). Empirical data indicated that Model A to be the most accurate representation of the demographic history (Supplementary Table 2). However, it is important to note that a low contribution (less than 5%) from an early OOA or outgroup of Eurasia could be misinterpreted as Model A (Supplementary Figures 6 and 7) but Model OX can be rejected with high certainty regardless of the admixture amount (Supplementary Figure 8).

We also ran models with low migration rates (m < 5 x 10^-5^, where m is the proportion of individuals moving from one population to another per generation) between modern human populations, and the results remained consistent (Supplementary Table 3). However, when we increased the migration rate (m < 5 x 10^-4^), although Model A remained the top-ranked model, it was no longer significantly better than Model O (Bayes factor between Model A and O [BF_AO_] <10, Supplementary Table 4). The higher migration rates lead to equifinality in the cSFS, which the neural network failed to distinguish^20,29^ and generally gave inconsistent results when running multiple times.

The best-fitting parameters for Model A (Table 1, Figure 2) largely correspond with the previously established OOA model, with some deviations specific to the inclusion of the PNG population^20,30–32^. Our model suggests that all OOA populations, including PNG, diverged from African populations (represented by Yoruba) around 62.4 (62 – 62.8) thousand years ago, experiencing a significant bottleneck (Table 1). Approximately 52 (51.6 – 52.8) thousand years ago, Neanderthals contributed around 3.7% (3.59 – 3.85%) of the genome to these OOA populations. Shortly thereafter, Europeans and East Asians diverged from the PNG populations around 51.2 (50.8 – 51.6) and 46.2 (45.9 – 46.5) thousand years ago, respectively. The PNG population then mixed with Denisovans around 31.2 (31.1 – 31.5) thousand years ago, contributing approximately 3.16% (3.05 – 3.21%) to the genome of PNG. Our analysis also shows that the PNG population experienced a more severe bottleneck (674 [663 – 689] of effective population size) than other OOA populations (i.e. Europeans 3512 [3423 – 3589] and East Asians 1771 [1730 – 1799] of effective population size), with growth rates significantly lower than those of other OOA populations, consistent with previously published data^6,33,34^. While our parameter inference is generally robust within the individual model, substantial changes occur when the underlying model is altered. Given that determining the precise demographic model for human populations is an ongoing effort, parameter estimates should be considered supplementary to the model rather than independent results.

**Figure 2:**
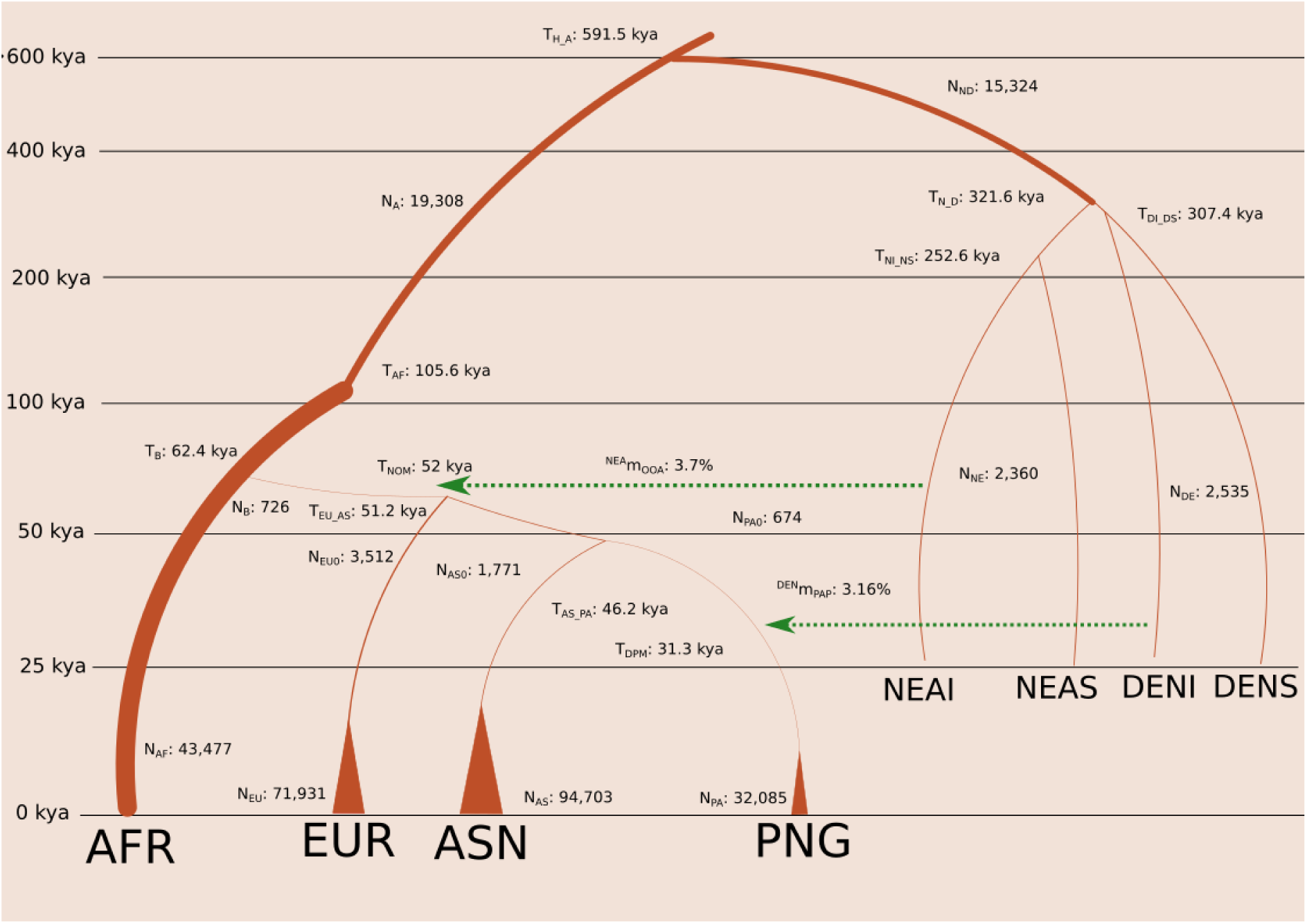
The schema of the Model A with the best fitted mean values of parameters coming from Table 1. AFR: Africa, EUR: European, ASN: East Asian, PAP: Papua New Guinean, NEAI: introgressed Neanderthal, NEAS: sequenced Altai Neanderthal, DENI: introgressed Denisova and DENS: sequenced Denisova. The corresponding values are written with the name of the parameters. The graph is not according to the scale. The y-axis represents time events from a thousand years ago (kya) from now.

**Table 1:** Best fitted Parameter values of Model A. CI: Confidence interval, ky: thousand years and kya: thousand years ago. PNG is Papua New Guinean and OOA is Out Of Africa.

| Params | Description | Mean (CI) | Events (kya) |
| --- | --- | --- | --- |
| $N_A$ | The ancestral effective population size | 19,308 (19,285 - 19,318) | |
| $N_{AF}$ | Effective population size of modern African population | 43,477 (43,048 - 43,957) | |
| $N_{EU}$ | Effective population size of modern European population | 71,931 (68,012 - 74,750) | |
| $N_{AS}$ | Effective population size of modern East Asian population | 94,703 (89,594 - 98,160) | |
| $N_{PA}$ | Effective population size of modern PNG population | 32,085 (30,889 - 33,303) | |
| $N_{NE}$ | Effective population size of Neanderthals | 2,360 (2,347 - 2,378) | |
| $N_{DE}$ | Effective population size of Denisova | 2,535 (2,519 - 2,549) | |
| $N_{EU0}$ | Effective population size of European population before exponential growth | 3,512 (3,423 - 3,589) | |
| $N_{AS0}$ | Effective population size of East Asian population before exponential growth | 1,771 (1,730 - 1,799) | |
| $N_{PA0}$ | Effective population size of PNG population before exponential growth | 674 (663 - 689) | |
| $N_B$ | Effective population size of OOA population | 726 (718 - 735) | |
| $N_{ND}$ | Effective population size of archaic hominin population | 15,324 (15,183 - 15,393) | |
| $T_{DPM}$ (ky) | Time interval of introgression of PNG population from Denisova | 31.3 (31.1 - 31.5) | 31.3 (31.1 - 31.5) |
| $T_{AS\_PA}$ (ky) | Time interval of separation between East Asian and PNG | 14.9 (14.6 - 15.1) | 46.2 (45.9 - 46.5) |
| $T_{EU\_AS}$ (ky) | Time interval of separation between East Asian and European | 5 (4.7 - 5.1) | 51.2 (50.8 - 51.6) |
| $T_{NOM}$ (ky) | Time interval of introgression of OOA population from Neanderthal | 0.9 (0.8 - 1) | 52 (51.6 - 52.8) |
| $T_B$ (ky) | Time interval of Separation of African and OOA population | 10.3 (10.2 - 10.4) | 62.4 (62 - 62.8) |
| $T_{AF}$ (ky) | Time interval of changes in effective population size of humans from ancestral population size | 43.2 (41.8 - 44.6) | 105.6 (104.1 - 107) |
| $T_{NI\_NS}$ (ky) | Time interval of Separation between sequenced Neanderthal and introgressed Neanderthal | 200.6 (198.6 - 203.3) | 252.6 (250.2 - 255.1) |
| $T_{DI\_DS}$ (ky) | Time interval of Separation between sequenced Denisoval and introgressed Denisova | 276 (274.6 - 277) | 307.4 (306.2 - 308.6) |
| $T_{N\_D}$ (ky) | Time interval of separation between Neanderthal and Denisova | 14.3 (12.8 - 15.5) | 321.6 (319.8 - 323.4) |
| $T_{H_A}$ (ky) | Time interval of separation of between modern humans and archaic hominins | 269.8 (268.8 - 271.4) | 591.5 (589.3 - 593.7) |
| $^{NEA}m_{OOA}$ (%) | Amount of Neanderthal introgression to OOA population | 3.7 (3.59 - 3.85) | |
| $^{DEN}m_{PAP}$ (%) | Amount of Denisova introgression to PNG population | 3.16 (3.05 - 3.21) | |

### Demographic model of PNG and RCCR curves

Relate analysis^21^ on simulated data coming from the best-fitting parameter values of Model A (Table 1) produced RCCR curves that replicated the empirical shift towards older divergence times for PNG versus Yoruba compared to other OOA populations (Figure 3). This shift appears to be primarily driven by the severe bottleneck and lower growth rate experienced by the PNG population (Supplementary Figure 9). Notably, a simpler model without any archaic introgression (with all other parameters remaining the same as in Table 1 except introgression) produced a similar shift (Supplementary Figure 10). When we simulated an African population with an earlier divergence, around 300 thousand years ago (similar to the San population)^35^, the RCCR shift became less pronounced as observed before^3,4^, indicating that the timing of African population divergence also influences the results (Supplementary Figure 11).

**Figure 3:**
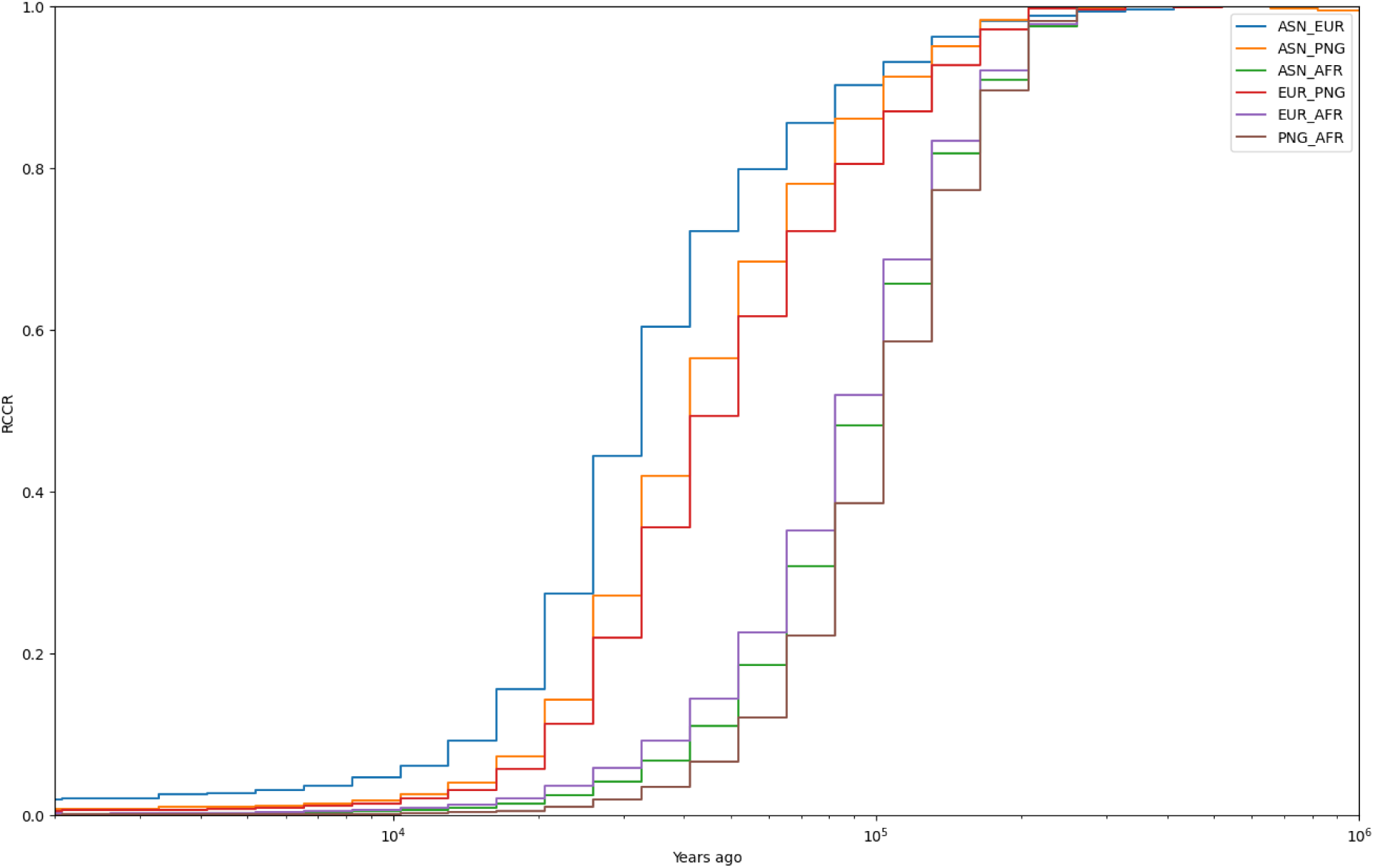
Relate results of relative cross coalescent curve on simulated data of best fitted values from Model A using 40 samples per population. AFR = African, EUR = European, ASN = East Asian and PNG = Papua New Guinean. The x-axis is years ago from modern times with the log scale. The y-axis is RCCR values calculated by Relate.

To further investigate this phenomenon, we conducted simulations of the PNG population under varying bottleneck intensities and growth rates using a minimal model to delimit the effects (see Methods for more details, Supplementary Table 5). The RCCR shift is influenced by both the bottleneck and the growth rate of the PNG population (Supplementary Figure 12). Further analysis of the most pronounced RCCR shift (produced from a bottleneck of 500 samples and no exponential growth for PNG) revealed that the estimates of coalescent rates for the PNG population, both within their population and cross population with Africa, are affected by this phenomenon. This in turn impacts the RCCR curve (Supplementary Figure 13), as RCCR is a ratio of these coalescent rate estimates [RCCR_01_= 2 x λ_01_ / (λ_0_ + λ_1_), where λ represents the coalescent rate, and 0 and 1 denote the two different populations]. The shift is primarily caused by changes in the coalescent rate estimates of PNG, which were affected due to the demographic history (changes in effective population size) occurring much before the actual event, extended over a much longer time period than the event itself^36^. Although the bottleneck of the PNG occurred around 50 thousand years ago, this event affected the coalescent rate estimate more than 100 thousand years ago. Thus the RCCR estimate deviated more for PNG than Europeans.

A similar pattern was observed in the RCCR curves between the PNG and other OOA populations (Figure 3). Although Europeans are the true outgroup of Asian and PNG populations in our simulated model, the RCCR analysis shows a greater separation between PNG and Europeans or East Asians than between Europeans and East Asians. This observation in empirical RCCR analysis was a key factor in the initial hypothesis that PNG are an outgroup to Europeans and East Asians^4^.

## Discussion

We successfully replicated the shift observed by Pagani et al.^4^, confirming its presence in both physically mapped and statistically phased sequences, which involved over 100 PNG samples. This consistency suggests that the shift is reproducible, though its underlying cause may differ from the original interpretation of Pagani et al.^4^. Our analysis using ABC-DLS supports a simpler demographic model for PNG populations, proposing them as a sister group to Asians with no substantial detectable contribution from an earlier OOA population. Interestingly, our simulated models reveal that a stronger bottleneck with a lower growth rate could produce a similar shift in RCCR analysis and potentially be misinterpreted as a signal of an earlier population separation. While RCCR is a valuable proxy for estimating the separation time between populations, it is not without biases. The shift could result from various factors, including earlier divergence times^22^, admixture with earlier diverged populations^4^, or even a bottleneck in one of the populations, as demonstrated in our study. Interestingly this demographic history of stronger bottleneck with slower growth rate was also experienced by the Andamanese population, which explains the shift found in the Andamanese population as well^37^. Thus, using RCCR analysis to rebuild the tree of divergence might need to be revised ^38,39^.

The observed shift in the RCCR curve suggests that a recent bottleneck can impact estimates of effective population size in the distant past. Notably, in our simulations, the Papua New Guinean bottleneck occurred much later (around 46.2 thousand years ago, as shown in Table 1) than the observed shift (peaking around 100 thousand years ago, as depicted in Figure 3) with a population (Yoruba) that separated a long time ago. This finding implies that the estimation of effective population size and cross-coalescent rates may not be entirely independent, potentially affecting RCCR analysis in its current form. Further analysis suggests (Supplementary Figure 12) that the estimation of coalescent rate was affected earlier than true changes of effective population size, which shifts the RCCR curve as RCCR is a ratio of coalescent rates. Additionally, this shift was absent in simulations involving populations that separated 300 thousand years ago, akin to the San population, indicating that the bottleneck effect diminishes over longer separation times.

Our results also reveal that when the contribution from an earlier OOA population is between 1-5%, our neural analysis misclassifies the Model AX to be Model A at a higher rate (Supplementary Figure 6). We found that when the contribution from an earlier OOA population is set between 1-5%, our ABC-DLS analysis tends to misclassify the Model AX as Model A at a higher rate A similar issue arises with Model M, where a low contribution (less than 5%) from an outgroup Eurasian population can still be misclassified as Model A (Supplementary Figure 7). Thus our analysis does not work for less than 5% contribution from these unknown ghost populations, though Model OX does not show a similar phenomenon with Model A misclassification (Supplementary Figure 8). While we cannot completely rule out the possibility of a small contribution from these populations, our analysis suggests that such models are not necessary to explain the RCCR shift as previously proposed ^4^.

Interestingly, our results position PNG as a sister group to Asian populations^11,12,18^ rather than an outgroup of European and Asian ^3,9,19^. The primary difference between those models and ours lies in the migration rates between populations. Previous models^20,29^ that incorporated significant migration rates between populations were found to have confounded results, leading us to avoid including migration rates in our models. Without migration, our Model O closely resembles the previous models of PNG ^3,9,19^. Given that those models used substantial migration rates^29^, they are not directly comparable to our models without migration rate. Indeed with high migration rates, our approach failed to distinguish between Model A and O with high certainty (Supplementary Table 4). Still our work suggests that the main lineage of PNG is coming from a sister group of Asia, which was not confounded by a convoluted migration rate patterns between populations.

Our parameter estimation suggests that the PNG population separated from other populations around 46.2 (45.9 – 46.5) thousand years ago, a timeline that aligns with archaeological estimates of when the ancestors of PNG reached the ancient continent of Sahul, the landmass that once connected New Guinea and Australia^1^. Additionally, our Relate analysis indicates that the separation time between PNG and European populations was the longest observed among OOA populations. However, as our model suggests, this is likely a bias caused by the bottleneck of PNG. This bottleneck may lead to an overestimation of the separation time, particularly in RCCR analysis. In reality, it is more likely that PNG and East Asian populations separated later than the divergence between PNG and European populations.

In conclusion, our study provides compelling evidence that the unique demographic events—specifically, a significant bottleneck and slower population growth—within the PNG population are key factors influencing the observed shifts in RCCR curves. These findings not only refine our understanding of PNG’s demographic history but also emphasise the necessity of accounting for population-specific demographic events when interpreting RCCR curves.

## Methods Relate

### Data

This study did not generate any new sequence data. For our analysis using Relate, we utilised a previously published Variant Calling Format (VCF) file^24^. This file was filtered and phased according to the methods outlined in the original publication. We phased the data using SHAPEIT v4.2^40^, opting for statistical phasing given the ample sample size. For further analysis, we retained only the Yoruba, British from England and Scotland, Han Chinese South^25,26^, both highlander and lowlander populations from PNG^24^ and Andamanese population^12^ (see Supplementary Table Table 7 for details).

After phasing, we only kept 40 random samples per population to make it equal (except for analysis of Andamanese in which case we used 10 samples per population). This random sampling process was repeated 10 times to calculate confidence intervals.

### Relate Analysis

We then used Relate_v1.1.8^21^ to generate genealogical trees and estimate the effective population size across all populations. Our analysis revealed that running the analysis on chromosome 1 alone produced results comparable to those obtained from the entire genome. Therefore, we limited our analysis to chromosome 1. To account for the impact of sample size on the results^21^, we maintained an equal sample size across populations. For cases with two or 10 samples per population (Supplementary Figures 4), we analysed all autosomal chromosomes to increase statistical power.

For the physically mapped data, we downloaded 10x Genomic Variant Calling Format (GVCF) data for two samples per population from the respective source^27^. The individual samples were merged using BCFtools merge^41^ and then analysed with Relate across all chromosomes (Supplementary Figure 3).

## ABC-DLS

### Data

All position VCF files were required before merging with archaic genomes to produce accurate Site Frequency Spectrum (SFS). Thus we downloaded GVCF files coming from from the high coverage 1000 genome dataset^25,26^ for the Yoruba (n=15) to represent African population, British from England and Scotland (n=15) to represent European population and Han Chinese South (n=15) to represent East Asian population. Additionally, we used fastq files from PNG highlanders (n=15) from our previous publication^24^. The samples were randomly selected from these datasets (see Supplementary Table 7). We also obtained all-position VCF files for Neanderthal from the Altai and Denisovan from their respective repositories^7,42^ and ancestral information was downloaded from the 1000 Genomes data^43^.

### Mapping

The fastq files from PNG samples were mapped to the GR38 reference genome using BWA-MEM v0.7.12^44^. The resulting BAM files were sorted and merged (for samples with multiple runs) using SAMtools v1.9^41,45^. Duplicates were marked using Picard tools from GATK v4.2.0.0^46^. and base recalibration was performed using BaseRecalibrator and ApplyBQSR, both from GATK. Finally, the BAM files were converted to CRAM format using SAMtools, adhering closely to GATK’s best practices ^47^.

### Variant Calling

We performed variant calling on the PNG CRAM files using the HaplotypeCaller from GATK to generate GVCF files. Since an unbiased SFS required all-position VCF files, we conducted variant calling using mpileup from BCFtools for all modern samples^48^.

### Liftover

Neanderthal and Denisovan VCF files, originally aligned to GR37, were lifted over to GR38 using LiftoverVcf from GATK^46^. We then merged the all-position VCF files from modern human genomes with those from archaic genomes using BCFtools. The resulting VCF files were used to create an unbiased SFS for both modern and archaic populations. This strategy helped us to retain variant sites that were mono-allelic or fixed in modern humans but not in archaic genomes. We filtered for biallelic SNPs present in every individual and annotated the VCF files with ancestral information using BCFtools query and annotate commands.

### Cross-population Site Frequency Spectrum

We filtered out positions with insufficient power for variant calling, following the strategy from our previous work^24^. We retained sites with a minimum alignment mapping quality and base quality of 20, and downgraded the mapping quality for reads with excessive mismatches using a coefficient of 50. Sites with depth coverage significantly higher or lower than the average across the dataset (by a factor of two) were also masked, based on criteria from our prior research^24^. Additional filtering was applied using the MSMC mappability mask^22^. Low-quality positions in Altai Neanderthal and Denisovan genomes were excluded, as were positions lacking ancestral allele information. This filtering left us with approximately 1.5 GB of regions with sufficient power for variant calling.

We found that Site Frequency Spectrum (SFS) quickly becomes intractable with higher numbers of samples per population. For instance, using five samples from each modern population, along with one Neanderthal and one Denisovan, produced 131,769 columns and 20,000 rows. Although neural networks can handle such large datasets, the process becomes computationally demanding, particularly for parameter estimation, which requires multiple ABC-DLS cycles to achieve precise estimates. To address this, we used cross-population SFS (cSFS) as a more efficient alternative to classical SFS. We generated two-dimensional SFS for each population pair, then flattened and concatenated these into a single-dimensional array. This approach reduced the amount of data to train significantly (e.g., from 131,769 to 999 columns more than 100 fold reduction), thereby improving the computational efficiency of neural network training.

We converted VCF files to SFS using the Run_VCF2SFS.py script, and then to cSFS using the Run_SFS_to_SFS2c.py script from ABC-DLS^20^. We selected five individuals from each modern human population (African, European, Asian, PNG), one Neanderthal, and one Denisovan sample. Multiple cSFS datasets were generated by varying the modern human samples while keeping the archaic samples constant. Ultimately, we obtained three cSFS datasets from 15 samples per modern human population and two archaics, which were concatenated row-wise. This multi-cSFS approach enhances parameter estimation by allowing median parameter values predicted from multiple cSFS datasets, thereby improving robustness in parameter estimation. Multiple cSFS is also useful for finding correct demographic models as the same top model produced by different cSFS suggest robustness of the model prediction.

### Simulation

We used msprime v1.2.0^49^ for simulating demographic models. These simulations generated cSFS datasets for training neural networks in ABC-DLS, and VCF files for input into Relate analysis.

We tested five demographic models (Supplementary Figure 5) based on the framework of Gravel et al.^30^, excluding migration rates between populations. All OOA populations experienced a significant bottleneck after diverging from African populations (Yoruba in this case). Neanderthals contributed genetic material to all OOA populations before the divergence of OOA populations^42^, while Denisovans contributed only to PNG^7^. Both introgressed archaic populations diverged from the sequenced archaic populations long before these events^9,50^. OOA populations then experienced exponential growth after the separation of Euroepans and East Asians^30^.

#### Model A

The PNG population is a sister group to Asians^11,12^.

#### Model O

The PNG population is an outgroup to European and Asian populations^3,19^.

#### Model M

A mixture of Models A and O, where the PNG population is a sister group to Asians but also carry genetic contributions from an unknown outgroup population of European and East Asians.

#### Model AX

Similar to Model M, but the unknown contributing population diverged from Africans before the main OOA event, resembling the first OOA hypothesis^4^.

#### Model OX

Similar to Model AX, but with the PNG population as an outgroup to European and Asian populations.

Priors for these models (Supplementary Table 1) were chosen from our empirical results and previous studies. Our strategy of ABC works as long as the true parameter values fall within the limit of priors. The priors do not have to be exact.

For instance, the modern effective population sizes were set as follows: European and East Asian populations ranged from 10,000 to 150,000, while African populations were set between 10,000 and 60,000, and PNG populations ranged from 10,000 to 50,000. This reflects the relatively modest population growth observed in the African and PNG populations in the Relate results, in contrast to the more pronounced population expansions in European and East Asian populations. The effective population sizes of the OOA populations prior to their exponential growth were set to much lower values, consistent with the well-documented bottleneck these populations experienced during the OOA event^6,30,51^, which is also supported by our Relate results. Neanderthal and Denisovan populations were assumed to have significantly smaller effective population sizes than modern humans, consistent with their known history of interbreeding^42,52^. The effective population size of the common ancestors of Neanderthals and Denisovans was set broadly, ranging from 500 to 30,000, due to the limited information available about this population.

We assumed that the archaic introgression into the PNG population occurred most recently, between 10,000 and 100,000 years ago (T_DPM_), based on the hypothesis that modern PNG population admixed with Denisovans after entering the Sunda and Sahul regions^53^. All other events in the model were assumed to occur consecutively before this timeframe, as detailed in Supplementary Table 6. For example, in Model A, the time interval of separation between PNG and East Asians is assumed to happen between 500 years and 50,000 years ago (T_EU_AS_), which corresponds to events ranging from 10,500 to 150,000 years ago in the simulation (T_DPM_ + T_EU_AS_). Whereas time intervals represent the parameter used in the simulation, events represent the timing of that event taking care of the past event that happened before that.

The level of introgression from archaic populations into modern humans was assumed to be between 1% and 5%, which is consistent with accepted estimates^42^. For the admixture from earlier OOA populations or outgroup Eurasian populations, we assumed a range of 1% to 99%, as analysis outside this range loses power to distinguish between models.

In certain scenarios, we incorporated migration rates between populations. Migration rates, denoted as m, represent the proportion of individuals moving from one population to another per generation, typically modelled as symmetric between populations. We applied these migration rates only to modern human populations (i.e., between Africans, Europeans, East Asians, and PNG) after their initial separation. The migration rates used in our models were m_ij_ < 5 x 10^-5^ and m_ij_ < 5 x 10^-4^, where i and j represent two different populations, and these values were used as priors.

#### Minimal model of OOA

To understand the effects of bottlenecks and exponential growth on RCCR curves, we simulated a minimal OOA model. African and OOA populations were set to have separated around 80 thousand years ago, each with an effective population size of 10,000 individuals. Europeans and PNG then diverged around 50 thousand years ago, with Europeans undergoing exponential growth (0.2% growth rate, leading to an effective population size of ∼300,000 in modern times). The PNG population experienced various combinations of bottleneck sizes (500 to 10,000 individuals) and growth rates (0% to 0.2%) (Supplementary Table 5).

### ABC and neural network analysis

We used ABC-DLS^20^ for model comparison and parameter estimation.

#### Model Comparison

We trained a neural network using 10,000 cSFS per model. For the simulations, we used 3 billion base pairs per sample, selecting five samples each from Yoruba, East Asian, European, and PNG populations, along with one sample from each archaic population (Altai-Neanderthal and Denisovan). The priors for the models are detailed in Table 1. The neural network predictions were then incorporated into the Approximate Bayesian Computation (ABC) framework to assess the confidence of these predictions.

#### Parameter Estimation

For parameter estimation, we trained the neural network with 20,000 cSFS in a single cycle. We began by running simulations with 100 MB (Mega Base) of sequence data per sample. As the simulation reached equilibrium, we progressively increased the sequence length to 500 MB, 1.5 GB, and finally 3 GB per sample, optimising the parameter estimates at each stage.

## Acknowledgements

This research was supported by the European Union through the European Regional Development Fund (Project No. 2014-2020.4.01.16-0030) to M.A.; and through Horizon 2020 research and innovation programme under grant no810645 and the European Regional Development Fund project no. MOBEC008 to A.E., M.A., M.Metspalu and M.Mondal. A.K.P. was supported by the Estonian Research Council grant PUT (PUTJD1186). M.Metspalu was supported by the Estonian Research Council grant PUT (PRG1899), the European Union’s Horizon Europe research and innovation programme under grant agreement No 101060011 (views and opinions expressed are however those of the author(s) only and do not necessarily reflect those of the European Union or European Research Executive Agency. Neither the European Union nor the granting authority can be held responsible for them) and the Estonian Research Council grant TK (TK214). F.-X.R.t and N. B were supported by the French Nationale Research Agency grant ANR-20-CE12-0003-01. Data analyses were carried out in part in the High-Performance Computing Center of the University of Tartu and supported by Tartu University grant no. PLTGI22904 to A.E.

## Author contributions

MMo, MMe and AE designed the study. MMo, MA and AKP performed data analysis. All the authors helped in writing the manuscript.

