## SupplementaryFile for "Resolving out of Africa event for Papua New Guinean population using neural network"

### Supplementary Figures

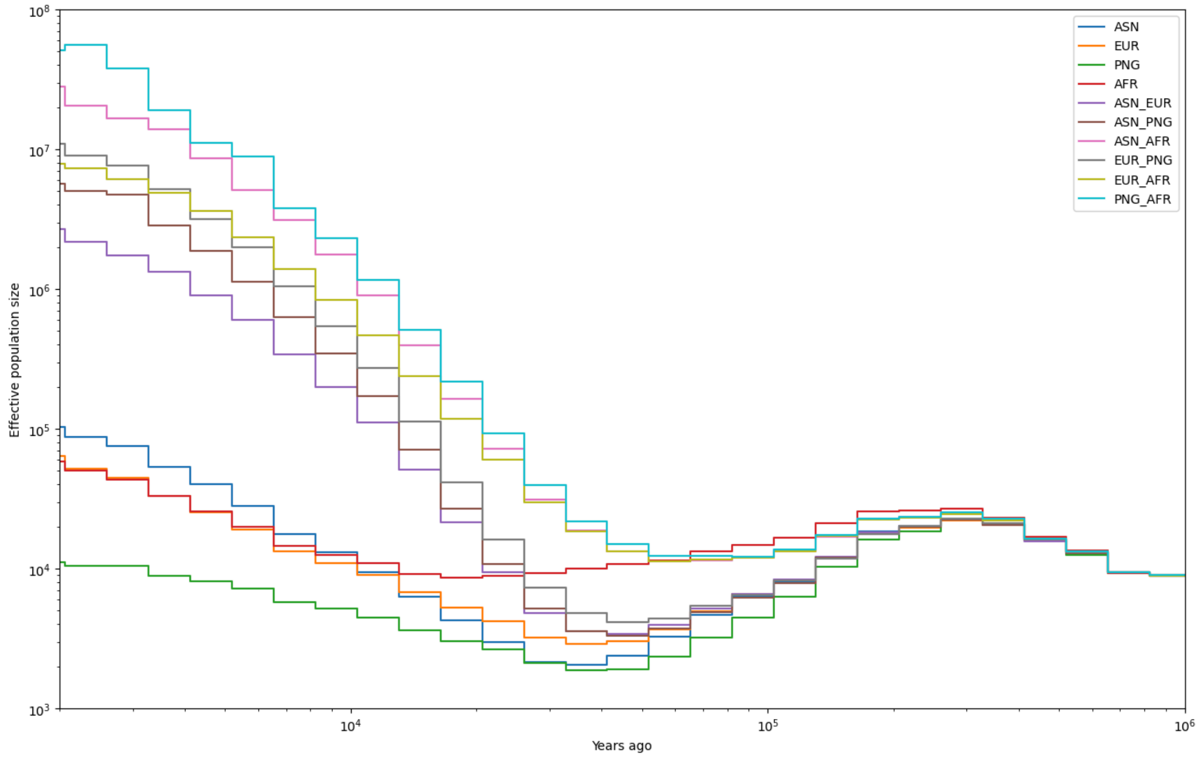


Supplementary Figure 1: Relate results of effective population size on empirical data using 40 samples per population. AFR = Yoruba, EUR = British from England and Scotland, ASN = Han Chinese and PNG = Highlanders of Papua New Guinea. The x-axis is years ago from modern times with the log scale. The y-axis is effective population size also in the log scale.

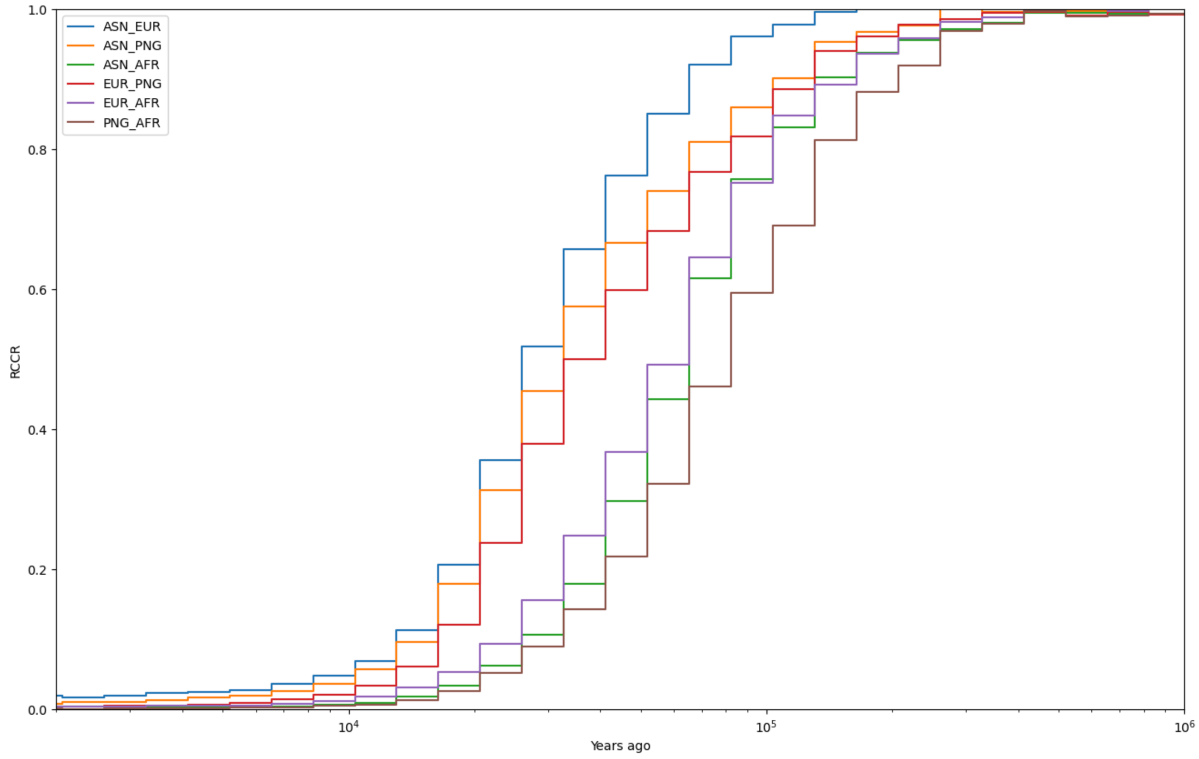


Supplementary Figure 2: Relate results of relative cross coalescent rate (RCCR) curve on empirical data using 40 samples per population. AFR = Yoruba, EUR = British from England and Scotland, ASN = Han Chinese and PNG = Lowlanders of Papua New Guinea. The x-axis is years ago from modern times with the log scale. The y-axis is RCCR values calculated by Relate.

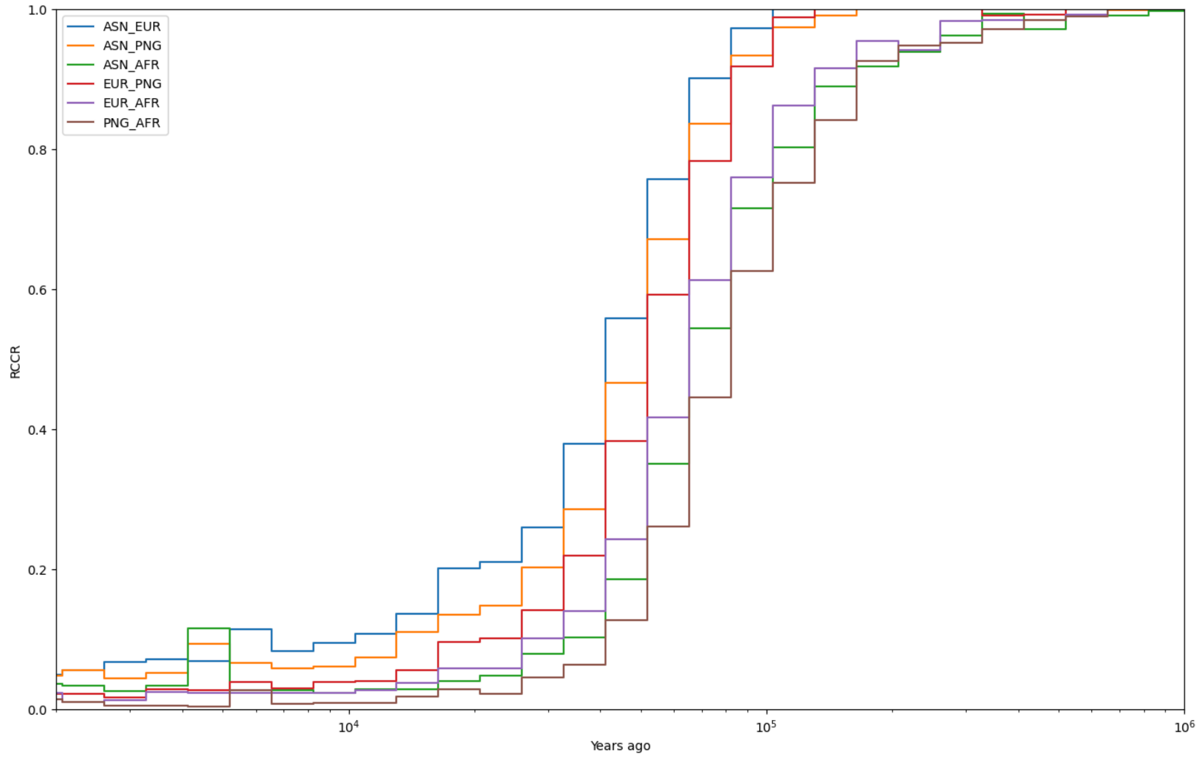


Supplementary Figure 3: Relate results of relative cross coalescent rate (RCCR) curve on physically mapped samples using two samples per population. AFR=Yoruba, EUR=Sardinian, ASN=Han and PNG=Highlands of Papua New Guinea. The x-axis is years ago from modern times with the log scale. The y-axis is RCCR values calculated by Relate.

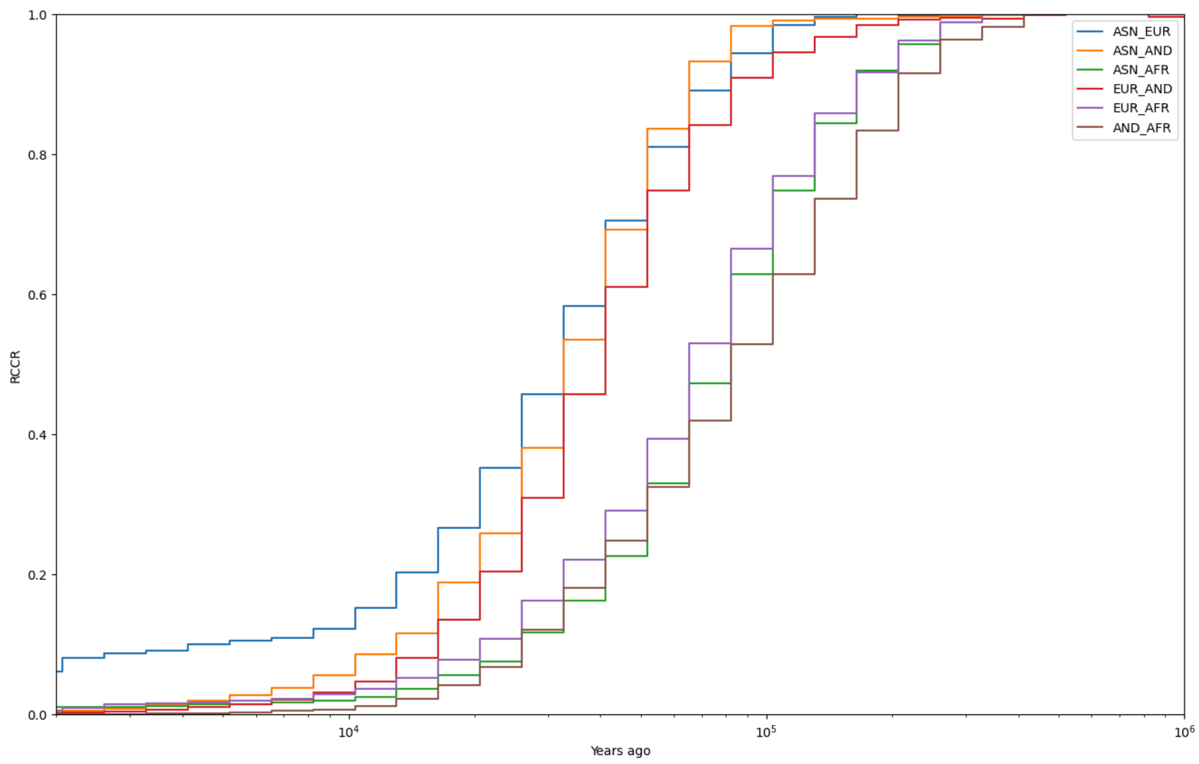


Supplementary Figure 4: Relate results of relative cross coalescent rate (RCCR) curve on empirical data using 10 samples per population. AFR = Yoruba, EUR = British from England and Scotland, ASN = Han Chinese and AND=Andamanese. The x-axis is years ago from modern times with the log scale. The y-axis is RCCR values calculated by Relate.

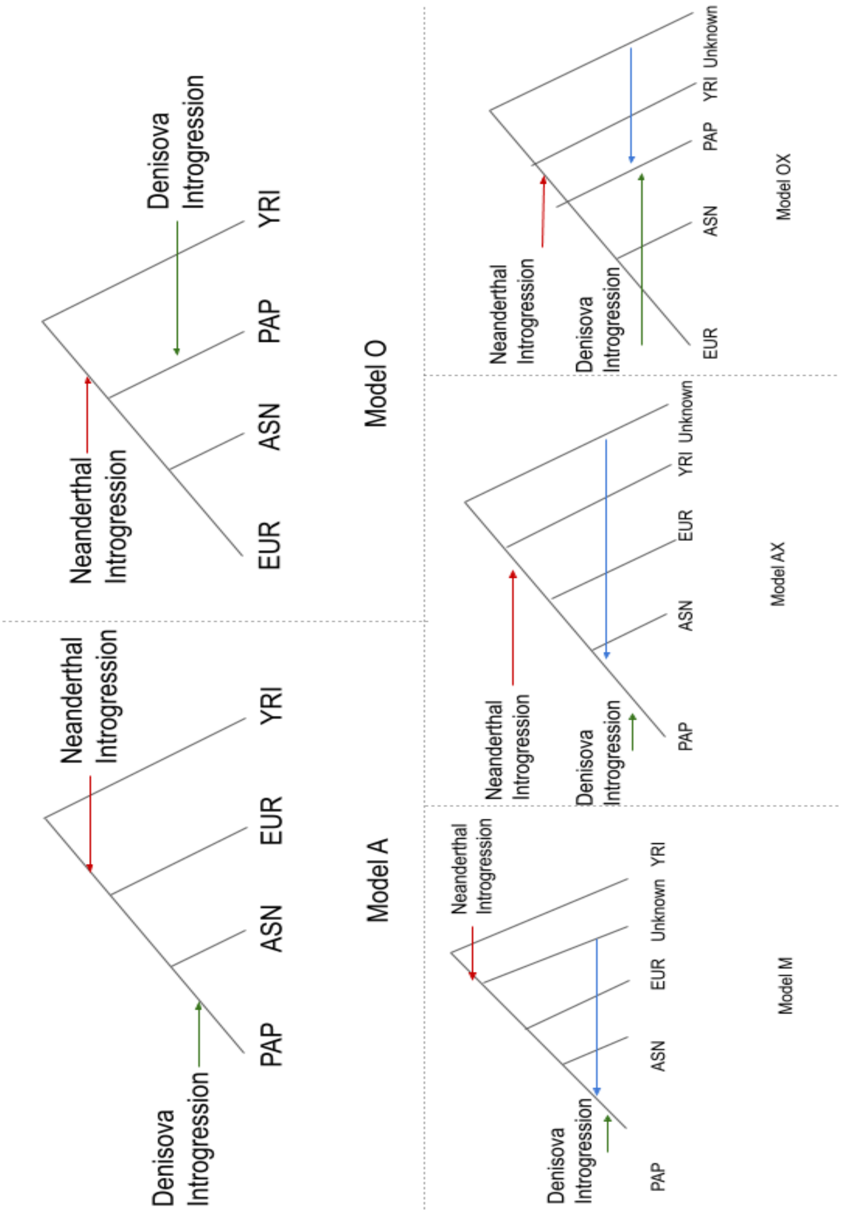


Supplementary Figure 5: The simplistic schema of the models tested

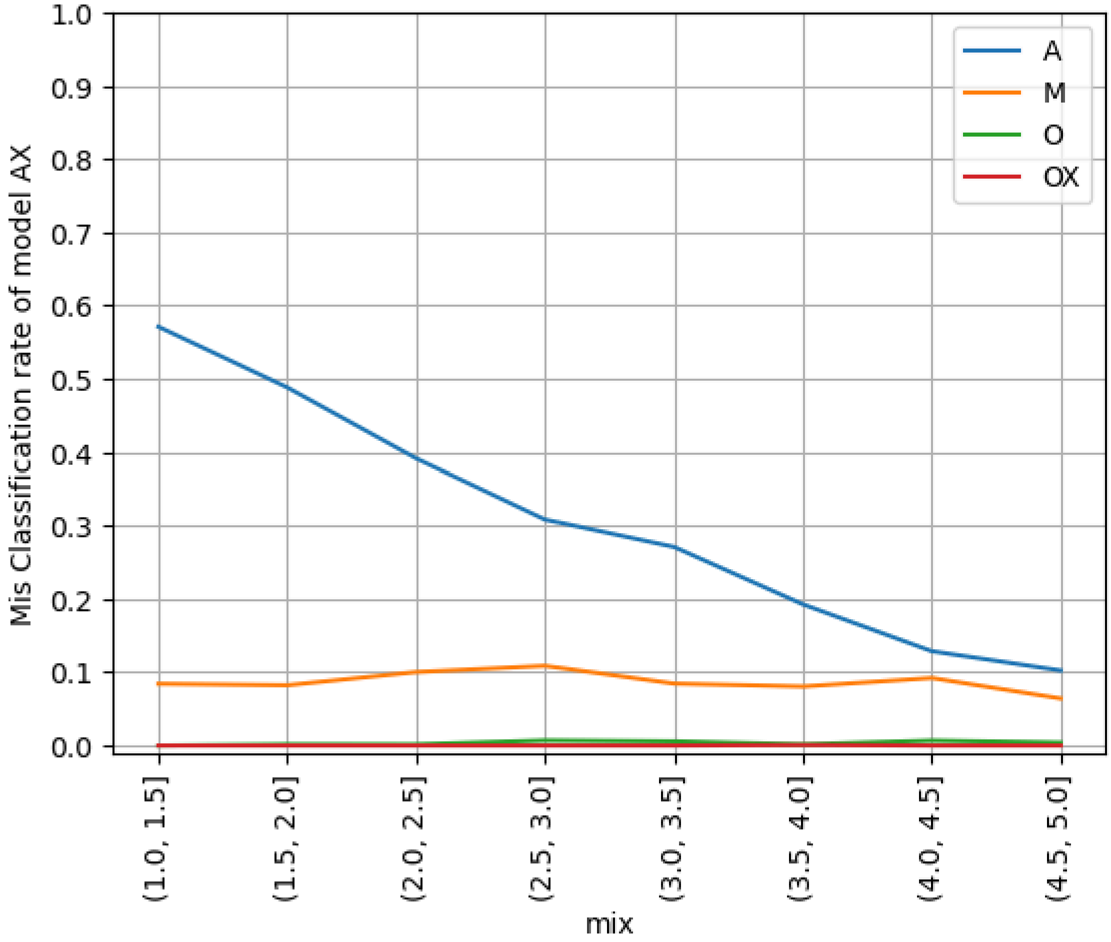


Supplementary Figure 6: Misclassification rate of ABC-DLS for Model AX to other model for admixture level less than 5% from earlier OOA population. The misclassification was calculated when ABC-DLS predicted the wrong model with a bayes factor more than 10.

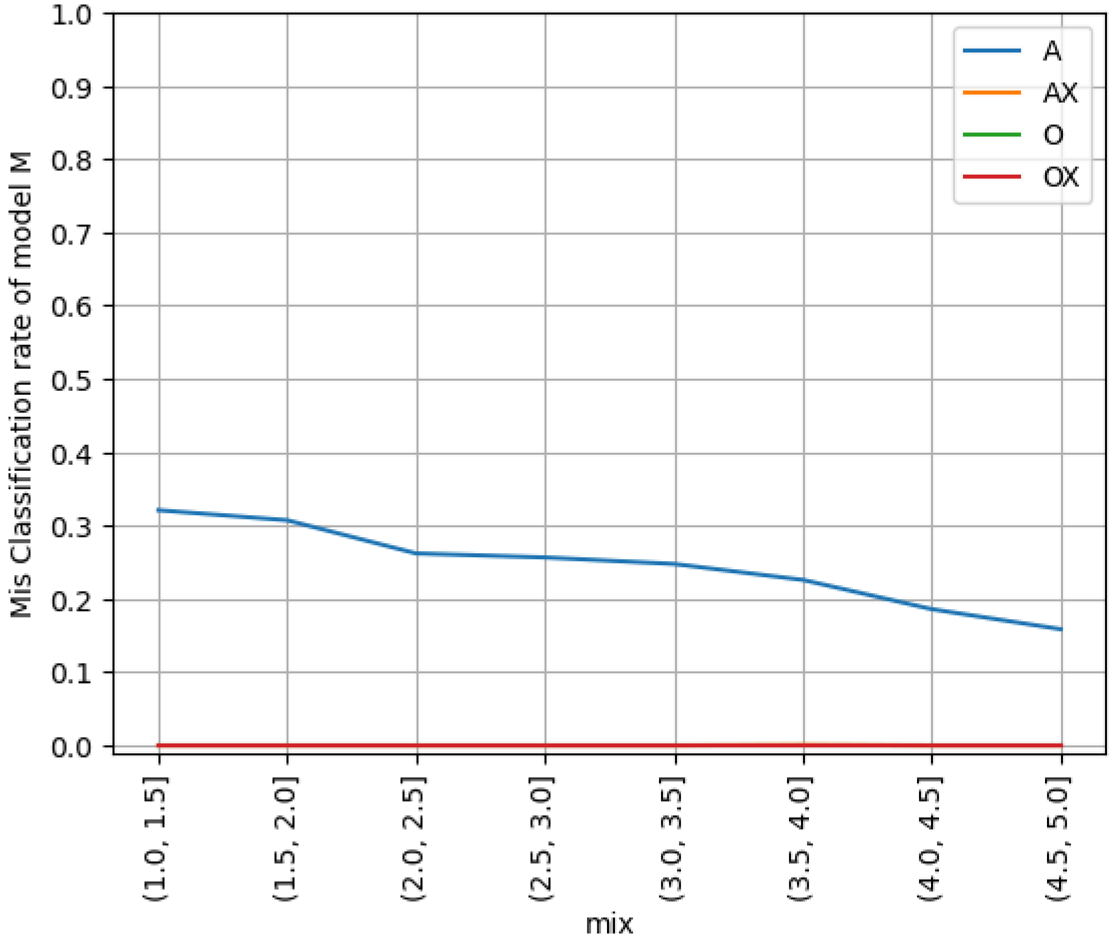


Supplementary Figure 7: Misclassification rate of ABC-DLS for Model M to other model for admixture level less than 5% from an outgroup of European and Asian. The misclassification was calculated when ABC-DLS predicted the wrong model with a bayes factor more than 10.

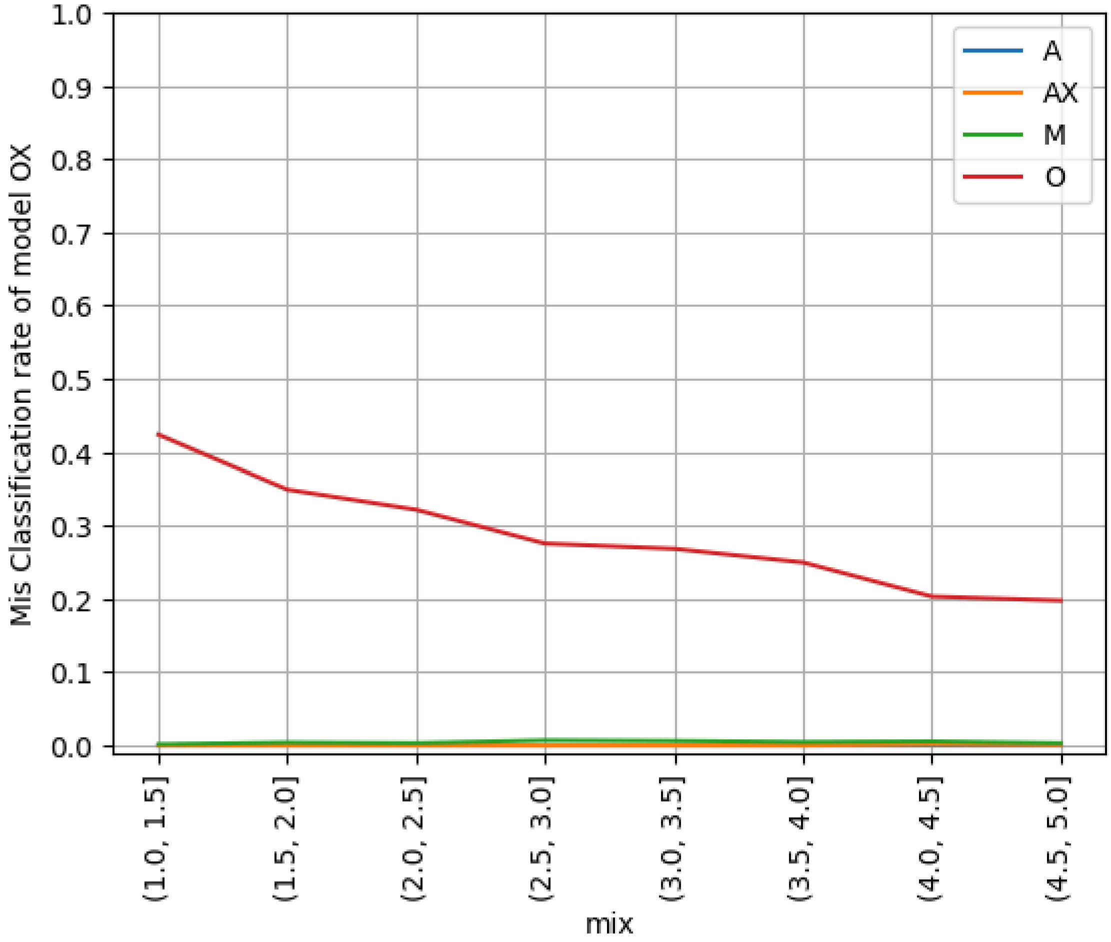


Supplementary Figure 8: Misclassification rate of ABC-DLS for Model OX to other model for admixture level less than 5% from earlier OOA population. The misclassification was calculated when ABC-DLS predicted the wrong model with a bayes factor more than 10.

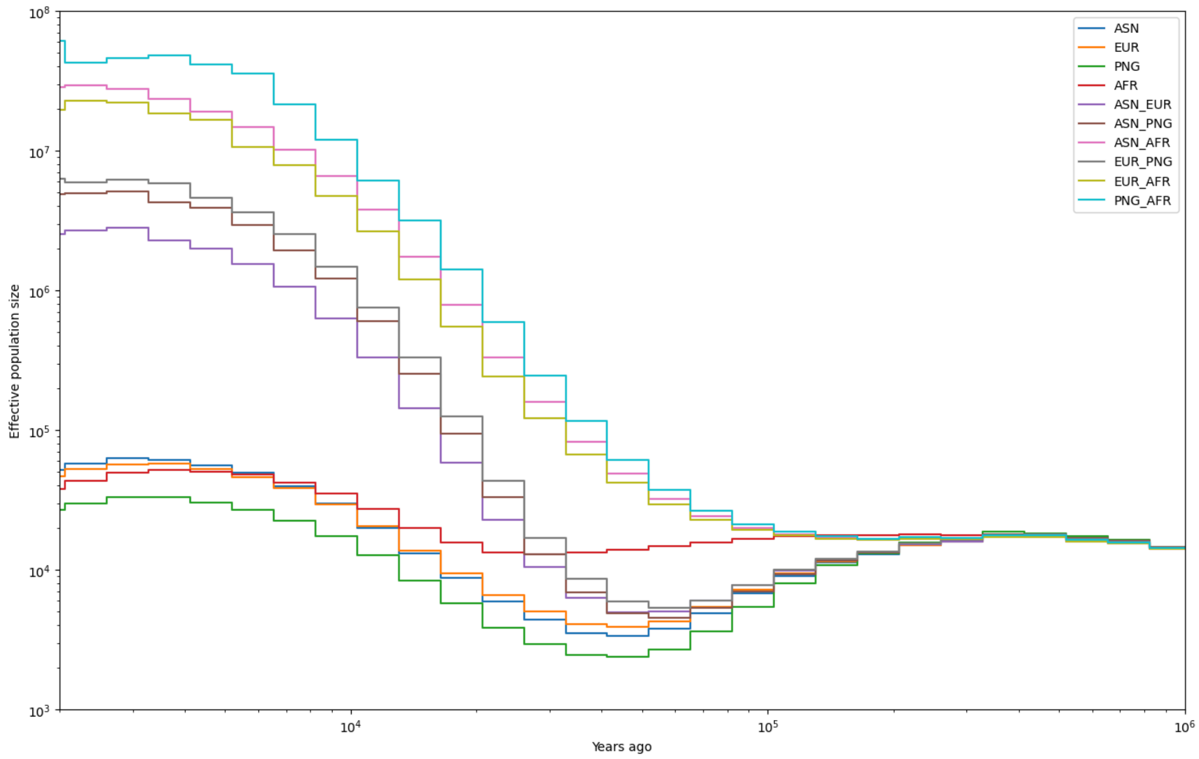


Supplementary Figure 9: Relate results of effective population size on simulated data of best fitted model A using 40 samples per population. AFR = Africa, EUR = European, ASN = East Asian and PNG = Papua New Guinea. The x-axis is years ago from modern times with the log scale. The y-axis is effective population size also in the log scale.

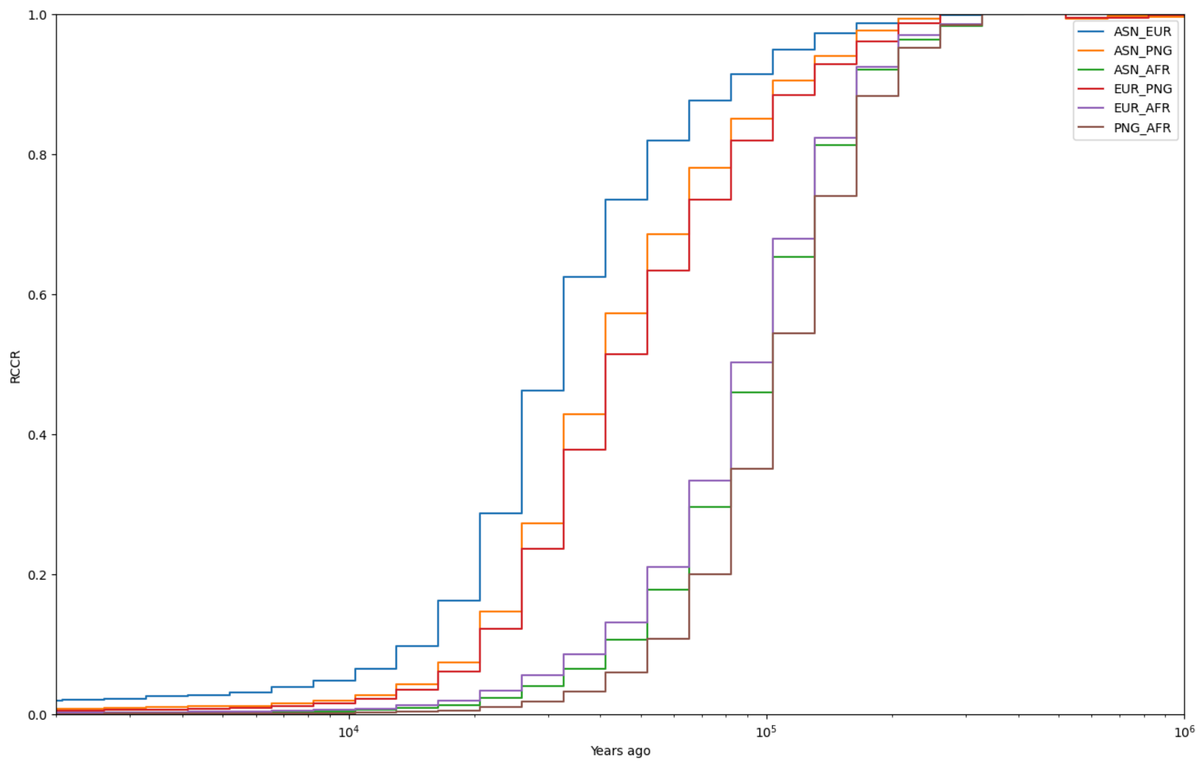


Supplementary Figure 10: Relate results of relative cross coalescent rate (RCCR) curve on simulated data of best fitted values from Model A without Archaic Introgression using 40 samples per population. AFR = African, EUR = European, ASN = East Asian and PNG = Papua New Guinean. The x-axis is years ago from modern times with the log scale. The y-axis is RCCR values calculated by Relate.

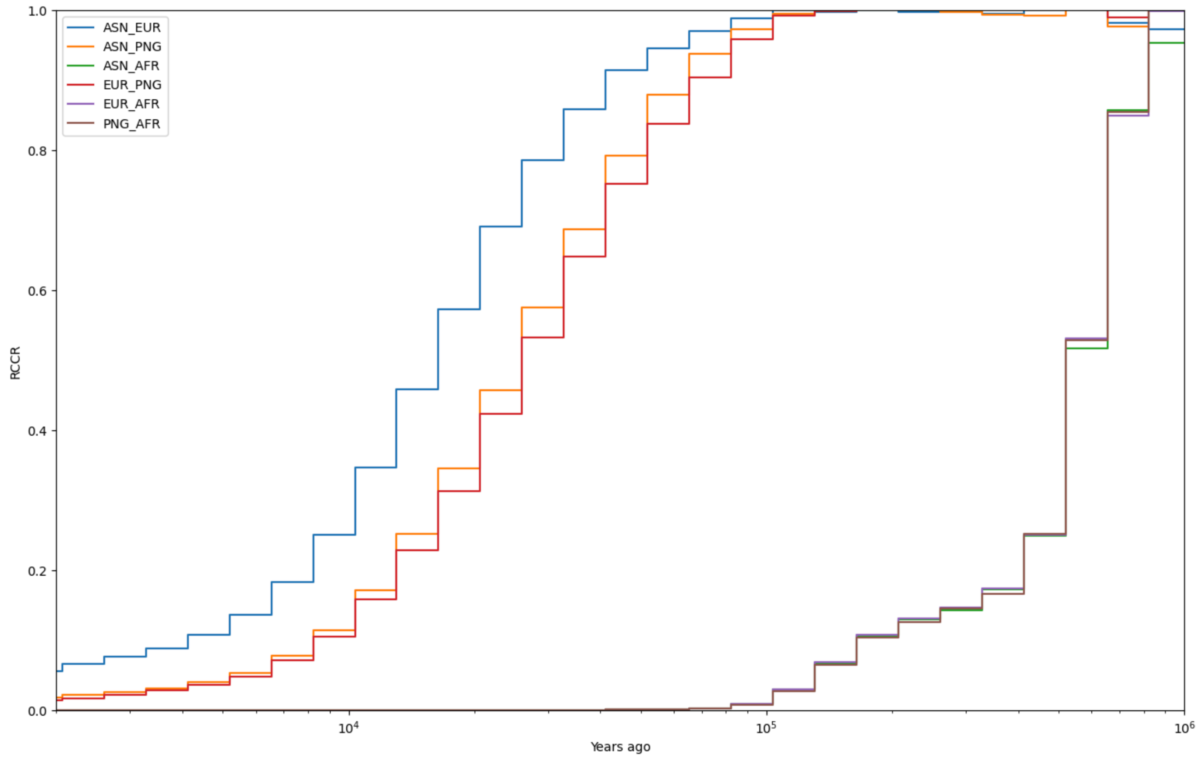


Supplementary Figure 11: Relate results of relative cross coalescent rate (RCCR) curve on simulated data of best fitted values from Model A with an African population separated 300 thousand years ago using 40 samples per population. AFR = African, EUR = European, ASN = East Asian and PNG = Papua New Guinean. The x-axis is years ago from modern times with the log scale. The y-axis is RCCR values calculated by Relate

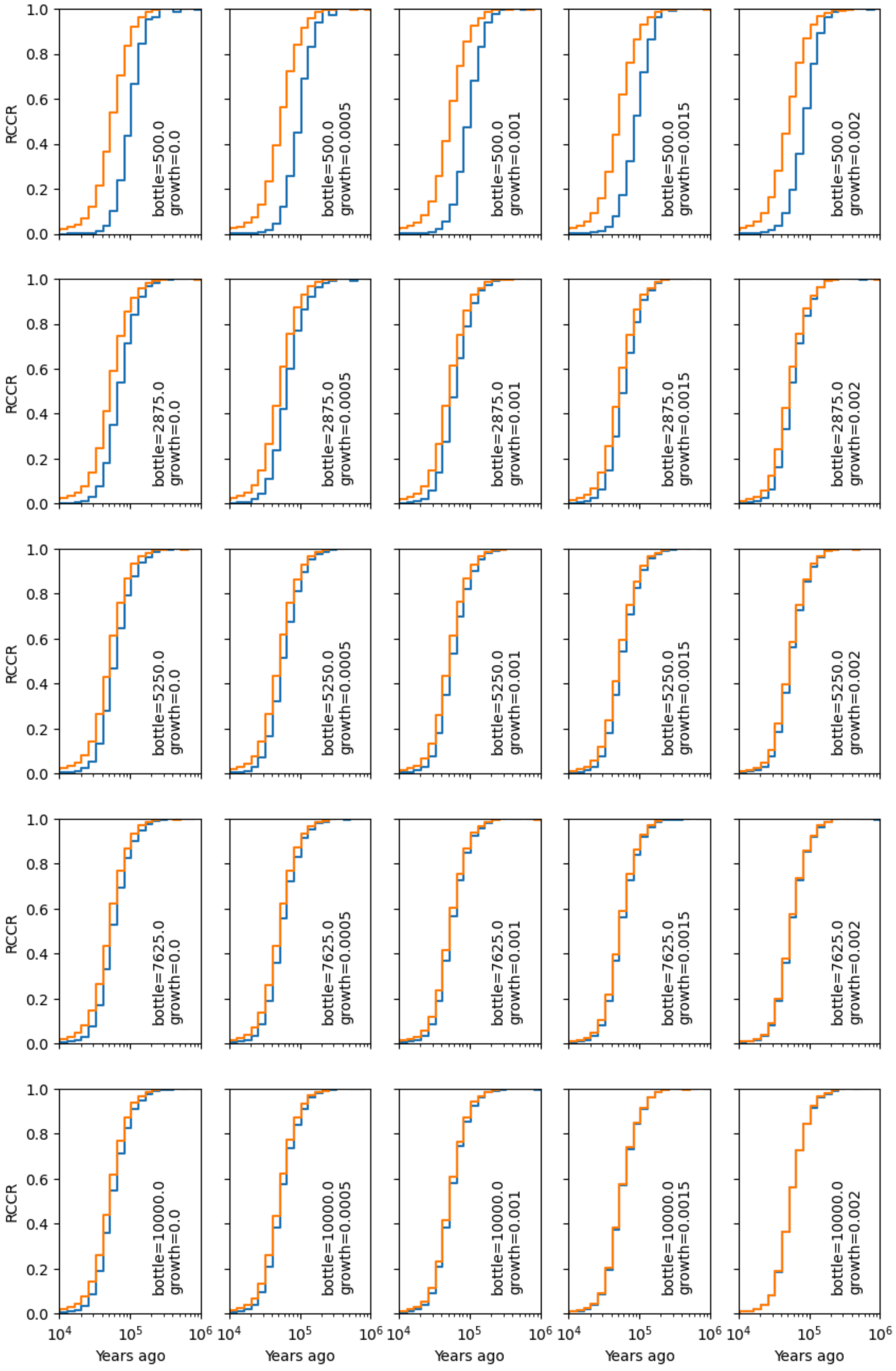
RCCR
Years ago
bottle=500.0 growth=0.0
bottle=500.0 growth=0.0005
bottle=500.0 growth=0.001
bottle=500.0 growth=0.0015
bottle=500.0 growth=0.002
bottle=2875.0 growth=0.0
bottle=2875.0 growth=0.0005
bottle=2875.0 growth=0.001
bottle=2875.0 growth=0.0015
bottle=2875.0 growth=0.002
bottle=5250.0 growth=0.0
bottle=5250.0 growth=0.0005
bottle=5250.0 growth=0.001
bottle=5250.0 growth=0.0015
bottle=5250.0 growth=0.002
bottle=7625.0 growth=0.0
bottle=7625.0 growth=0.0005
bottle=7625.0 growth=0.001
bottle=7625.0 growth=0.0015
bottle=7625.0 growth=0.002
bottle=10000.0 growth=0.0
bottle=10000.0 growth=0.0005
bottle=10000.0 growth=0.001
bottle=10000.0 growth=0.0015
bottle=10000.0 growth=0.002

Supplementary Figure 12: RCCR graph with combination of bottleneck and growth rate using simulated model using Minimal model of the out of Africa. Orange line is the Africa vs European RCCR curve which is constant throughout the graph, whereas blue line is the Africa vs Papua New Guinean RCCR curve under different amount of bottleneck and growth rate (written inside the legend) for Papua New Guinean population.

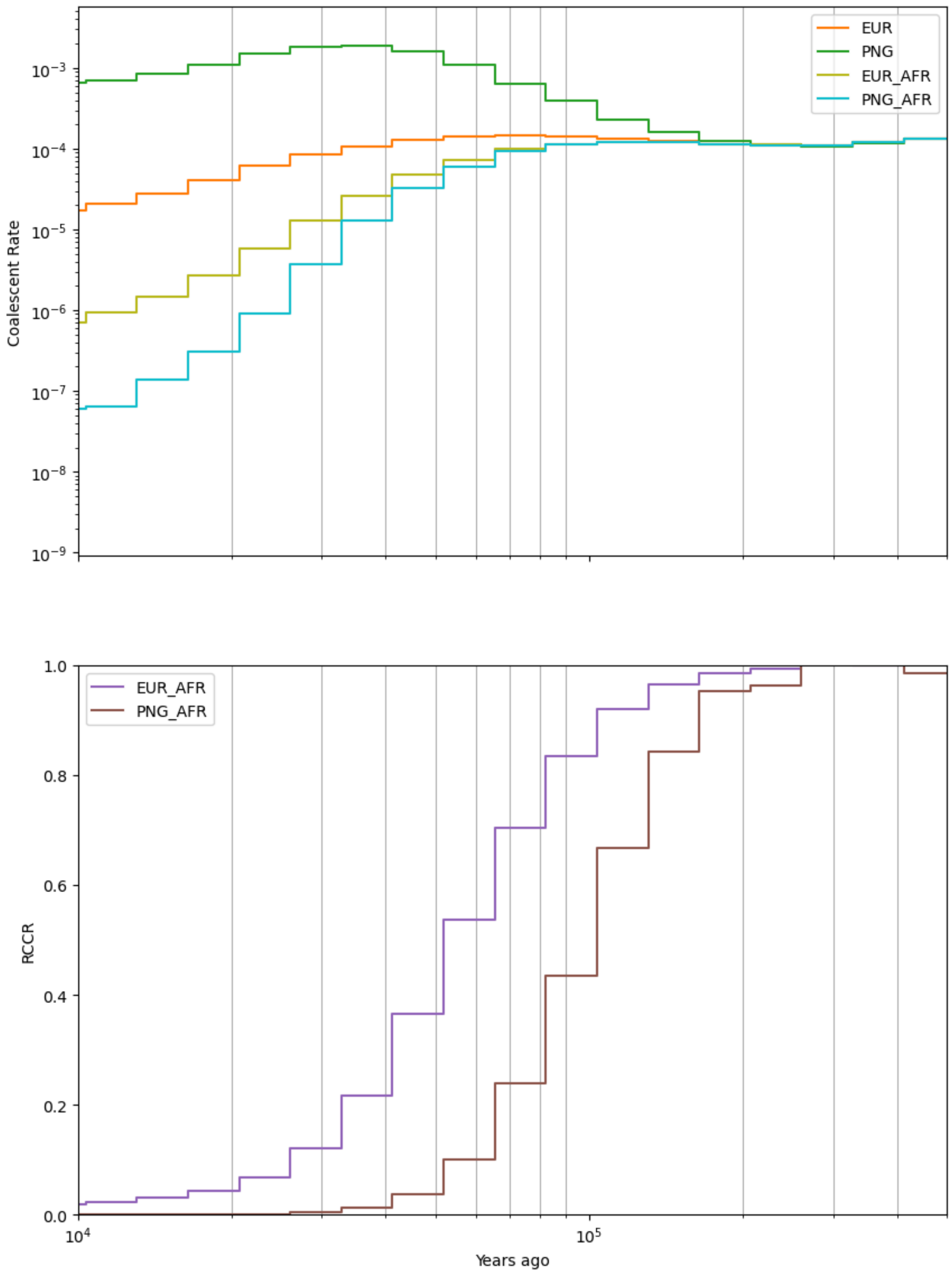

EUR
PNG
EUR_AFR
PNG_AFR
Coalescent Rate
$10^{-3}$
$10^{-4}$
$10^{-5}$
$10^{-6}$
$10^{-7}$
$10^{-8}$
$10^{-9}$
EUR_AFR
PNG_AFR
RCCR
1.0
0.8
0.6
0.4
0.2
0.0
$10^{4}$
$10^{5}$
Years ago

Supplementary Figure 13: Coalescent rate and corresponding RCCR curve of a simple simulated model for the top separated RCCR curve between Africa vs European and Africa vs Papua New Guinean. The Papua New Guinean has a bottleneck of 500 individuals and growth rate 0. The European population never had a bottleneck and growth rate of 0.2%. EUR = European and PNG = Papua New Guinean.

### Supplementary Tables

| Params | Model A | Model O | Model M | Model AX | Model OX |
|---|---|---|---|---|---|
| $N_A$ | 5,000 - 25,000 | 5,000 - 25,000 | 5,000 - 25,000 | 5,000 - 25,000 | 5,000 - 25,000 |
| $N_{AF}$ | 10,000 - 60,000 | 10,000 - 60,000 | 10,000 - 60,000 | 10,000 - 60,000 | 10,000 - 60,000 |
| $N_{EU}$ | 10,000 - 150,000 | 10,000 - 150,000 | 10,000 - 150,000 | 10,000 - 150,000 | 10,000 - 150,000 |
| $N_{AS}$ | 10,000 - 150,000 | 10,000 - 150,000 | 10,000 - 150,000 | 10,000 - 150,000 | 10,000 - 150,000 |
| $N_{PA}$ | 5,000 - 50,000 | 5,000 - 50,000 | 5,000 - 50,000 | 5,000 - 50,000 | 5,000 - 50,000 |
| $N_{NE}$ | 500 - 10,000 | 500 - 10,000 | 500 - 10,000 | 500 - 10,000 | 500 - 10,000 |
| $N_{DE}$ | 500 - 10,000 | 500 - 10,000 | 500 - 10,000 | 500 - 10,000 | 500 - 10,000 |
| $N_{EU0}$ | 500 - 5,000 | 500 - 5,000 | 500 - 5,000 | 500 - 5,000 | 500 - 5,000 |
| $N_{AS0}$ | 500 - 5,000 | 500 - 5,000 | 500 - 5,000 | 500 - 5,000 | 500 - 5,000 |
| $N_{PA0}$ | 500 - 5,000 | 500 - 5,000 | 500 - 5,000 | 500 - 5,000 | 500 - 5,000 |
| $N_B$ | 500 - 5,000 | 500 - 5,000 | 500 - 5,000 | 500 - 5,000 | 500 - 5,000 |
| $N_{ND}$ | 500 - 30,000 | 500 - 30,000 | 500 - 30,000 | 500 - 30,000 | 500 - 30,000 |
| $T_{DPM}$ (ky) | 10 - 100 | 10 - 100 | 10 - 100 | 10 - 100 | 10 - 100 |
| $T_{AS\_PA}$ (ky) | 0.5 - 50 | NA | NA | NA | NA |
| $T_{Mix}$ (ky) | NA | NA | 0.5 - 50 | 10 - 100 | 0.5 - 50 |
| $T_{EU\_AS}$ (ky) | 0.5 - 50 | 10 - 100 | 0.5 - 50 | 0.5 - 50 | 10 - 100 |
| $T_{EA\_PA}$ (ky) | NA | 0.5 - 50 | 0.5 - 50 | NA | 0.5 - 50 |
| $T_{NOM}$ (ky) | 0.5 - 50 | 0.5 - 50 | 0.5 - 50 | 0.5 - 50 | 0.5 - 50 |
| $T_B$ (ky) | 0.5 - 50 | 0.5 - 50 | 0.5 - 50 | 0.5 - 50 | 0.5 - 50 |
| $T_{X\_H}$ (ky) | NA | NA | NA | 0.5 - 100 | 0.5 - 100 |
| $T_{AF}$ (ky) | 0.5 - 200 | 0.5 - 200 | 0.5 - 200 | 0.5 - 200 | 0.5 - 200 |
| $T_{NI\_NS}$ (ky) | 10 - 500 | 10 - 500 | 10 - 500 | 10 - 500 | 10 - 500 |
| $T_{DI\_DS}$ (ky) | 10 - 500 | 10 - 500 | 10 - 500 | 10 - 500 | 10 - 500 |
| $T_{N\_D}$ (ky) | 10 - 500 | 10 - 500 | 10 - 500 | 10 - 500 | 10 - 500 |
| $T_{H\_A}$ (ky) | 100 - 500 | 100 - 500 | 100 - 500 | 100 - 500 | 100 - 500 |
| ${}^{NEA}m_{OOA}$ (%) | 1 - 5 | 1 - 5 | 1 - 5 | 1 - 5 | 1 - 5 |
| ${}^{DEN}m_{PAP}$ (%) | 1 - 5 | 1 - 5 | 1 - 5 | 1 - 5 | 1 - 5 |
| Mix (%) | NA | NA | 1 - 99 | 1 - 99 | 1 - 99 |

Supplementary Table 1: Prior range for different models; ky = thousand years

| | A | O | M | AX | OX |
|---|---|---|---|---|---|
| A | 84.89% | 0.04% | 12.33% | 2.74% | 0.00% |
| O | 0.33% | 81.97% | 6.25% | 0.42% | 11.03% |
| M | 13.95% | 5.86% | 71.86% | 7.00% | 1.33% |
| AX | 6.21% | 0.13% | 10.37% | 80.17% | 3.12% |
| OX | 0.00% | 11.28% | 0.78% | 7.08% | 80.87% |
| Empirical | 94.96% | 0.00% | 1.93% | 3.12% | 0.00% |

Supplementary Table 2: ABC-DLS classification results

| | A | O | M | AX | OX |
|---|---|---|---|---|---|
| A | 87.34% | 0.71% | 7.95% | 4.00% | 0.00% |
| O | 1.08% | 87.41% | 9.95% | 1.48% | 0.07% |
| M | 9.51% | 8.07% | 62.38% | 19.61% | 0.44% |
| AX | 2.12% | 1.23% | 22.24% | 74.29% | 0.12% |
| OX | 0.00% | 0.00% | 0.00% | 0.00% | 100.00% |
| Empirical | 92.45% | 0.00% | 3.76% | 3.79% | 0.00% |

Supplementary Table 3: ABC-DLS results on classification with low migration rate ($m < 5 \times 10^{-5}$)

| | A | O | M | AX | OX |
|---|---|---|---|---|---|
| A | 62.25% | 16.28% | 15.63% | 5.81% | 0.02% |
| O | 14.69% | 63.50% | 12.08% | 9.68% | 0.05% |
| M | 14.70% | 10.96% | 55.74% | 18.59% | 0.00% |
| AX | 9.03% | 8.01% | 21.94% | 61.01% | 0.00% |
| OX | 0.00% | 0.00% | 0.00% | 0.00% | 100.00% |
| Empirical | 48.37% | 44.33% | 4.85% | 2.45% | 0.00% |

Supplementary Table 4: ABC-DLS results on classification with high migration rate ($m < 5 \times 10^{-4}$)

| Params | Description | Values |
|---|---|---|
| $N_{AF}$ | Effective population size of modern African population | 10,000 |
| $N_{EU0}$ | Effective population size of European population before exponential growth | 10,000 |
| $r_{EU}$ | Increase of effective population size for Europeans per generation for exponential growth | 0.2 % |
| $N_{PA0}$ | Effective population size of PNG population before exponential growth | 500-10,000 |
| $r_{PA}$ | Increase of effective population size for PNG per generation for exponential growth | 0-0.2 % |
| $T_{EU\_PA}$ | Time event of separation between Europeans and PNG | 50 kya |
| $T_B$ | Time event of Separation of African and out of Africa population | 80 kya |

Supplementary Table 5: Parameter values used for a Minimal model of OOA

| Params | Model A | Model O | Model M | Model AX | Model OX |
|---|---|---|---|---|---|
| **Denisova introgression to PNG** | $T_{DPM}$ | $T_{DPM}$ | $T_{DPM}$ | $T_{DPM}$ | $T_{DPM}$ |
| **Separation between PNG and East Asian (A_P)** | $T_{DPM} + T_{AS\_PA}$ | NA | NA | NA | NA |
| **Admixture between Asian and outgroup PNG (A_O)** | NA | NA | $T_{DPM} + T_{Mix}$ | NA | NA |
| **Admixture between Asian and earlier OOA (A_X)** | NA | NA | NA | $T_{DPM}+ T_{Mix}$ | NA |
| **Admixture between outgroup PNG and earlier OOA (O_X)** | NA | NA | NA | NA | $T_{DPM}+ T_{Mix}$ |
| **Separation between European and Asian (E_A)** | A_P + $T_{EU\_AS}$ | $T_{EU\_AS}$ | A_O + $T_{EU\_AS}$ | AX + $T_{EU\_AS}$ | $T_{EU\_AS}$ |
| **Separation between Eurasian and outgroup PNG (EA_PA)** | NA | max ($T_{DPM}$ + $T_{EU\_AS}$) + $T_{EA\_PA}$ | E_A + $T_{EA\_PA}$ | NA | max (E_A, O_X) + $T_{EA\_PA}$ |
| **Neanderthal introgression to OOA population (NI)** | E_A + $T_{NOM}$ | EA_PA + $T_{NOM}$ | EA_PA + $T_{NOM}$ | EA_PA + $T_{NOM}$ | EA_PA + $T_{NOM}$ |
| **Separation between OOA and** | NI + $T_B$ | NI + $T_B$ | NI + $T_B$ | NI + $T_B$ | NI + $T_B$ |

| | | | | | |
|---|---|---|---|---|---|
| **African (OOA)** | | | | | |
| **Separation between earlier OOA and African (xOOA)** | NA | NA | NA | OOA + $T_{X\_H}$ | OOA + $T_{X\_H}$ |
| **Ancestral Size change** | OOA + T_AF | OOA + $T_{AF}$ | OOA + $T_{AF}$ | xOOA+ $T_{AF}$ | xOOA+ $T_{AF}$ |
| **Separation between Introgressed Neanderthal and Sequenced Neanderthal (NI_NS)** | NI + $T_{NI\_NS}$ | NI + $T_{NI\_NS}$ | NI + $T_{NI\_NS}$ | NI + $T_{NI\_NS}$ | NI + $T_{NI\_NS}$ |
| **Separation between Introgressed Denisova and Sequenced Denisova (DI_DS)** | $T_{DPM}$ + $T_{DI\_DS}$ | $T_{DPM}$ + $T_{DI\_DS}$ | $T_{DPM}$ + $T_{DI\_DS}$ | $T_{DPM}$ + $T_{DI\_DS}$ | $T_{DPM}$ + $T_{DI\_DS}$ |
| **Split between Neanderthal Denisova (N_D)** | max(NI_NS, DI_DS) + $T_{N\_D}$ | max(NI_NS, DI_DS) + $T_{N\_D}$ | max(NI_NS, DI_DS) + $T_{N\_D}$ | max(NI_NS, DI_DS) + $T_{N\_D}$ | max(NI_NS, DI_DS) + $T_{N\_D}$ |
| **Split Between Human and Archaics** | max (N_D, OOA) + $T_{H\_A}$ | max (N_D, OOA) + $T_{H\_A}$ | max (N_D, OOA) + $T_{H\_A}$ | max (N_D, xOOA) + $T_{H\_A}$ | max (N_D, xOOA) + $T_{H\_A}$ |

Supplementary Table 6: Relations of events with the time intervals used in the simulations
